# Regional reconfiguration of functional brain networks during childhood and adolescence: evaluating age and sex effect

**DOI:** 10.64898/2026.05.21.726818

**Authors:** Catherine Zhiqian Fang, Hajer Nakua, Xin Ma, Aiying Zhang, Seonjoo Lee

## Abstract

**Introduction:** While global topological properties of brain networks reach relative maturity early in development, functional reconfigurations at the regional level continue throughout adolescence to support cognitive maturation. However, regional age and sex-specific developmental patterns of functional reconfiguration remain incompletely understood.

**Methods:** We analyzed resting-state fMRI data from 528 participants aged 5–21 years from the Human Connectome Project in Development. Three regional graph-theory metrics (betweenness centrality, hub score, and local efficiency) were computed for each individual’s functional network. Cognition was measured using NIH toolbox. Parallel factor analysis was employed to decompose an individual × region × metric array into factors representing distinct developmental properties in the full sample and separately for males and females. Brain-cognition associations were examined in developmental subgroups (<13, 13-18, >18 years).

**Results:** Three factors emerged, characterizing visual, multimodal integration, and higher-order factors. Across development, metrics capturing network integration (betweenness centrality and hubness) showed general stability, while metrics capturing segregation (local efficiency) presented distinct peaks, particularly in the visual factor. Females showed earlier peaks and declines in higher-order factor, while males exhibited greater variability and protracted maturation in multimodal and higher-order factors. Brain-cognition associations were modest with early childhood and crystallized cognition composites showed small negative correlations with hub score in entire sample (r=-0.212) and local efficiency in males aged <13 years (r=-0.215).

**Conclusion:** Findings highlight nonlinear, sex-specific functional reconfiguration at region-level during childhood and adolescence, underscoring the importance of sex-stratified analyses in developmental and providing a crucial foundation for future investigations of developmental disorders.

## 1. Introduction

Throughout childhood and adolescence, functional brain networks reorganize extensively to form a mature functional architecture to support increasingly complex cognitive processing (Menon, 2013). Global brain architecture, including network affiliation of each brain region, is largely established early in development (Gao et al., 2015), but refinements within and between networks continue to unfold during adolescence, giving rise to variation in cognitive development (Luna & Sweeney, 2004). Brain regions communicate both within and between networks. Between-network connections increase across development (Grayson & Fair, 2017), supporting greater network integration and more efficient communication across distributed regions (Fair et al., 2009). During development, brain connectivity shifts from primarily short-range connections to more extensive long-range ones (Dosenbach et al., 2010; Fair et al., 2009; Supekar et al., 2009). This transition leads to clearer definitions of network functions and increased integration among brain regions as age increases (Uddin et al., 2011). Specifically, two distinct developmental patterns emerge during adolescence. Regions specialized in somatosensory and motor functions continue to strengthen their existing connections, whereas regions involved in complex cognitive processes (e.g., subcortical regions) undergo remodeling such that connections that were weak during childhood strengthen, while those strong connections weaken with age (Hettwer et al., 2024; Váša et al., 2020). Such reconfiguration underpins the emergence of complex cognitive functions, such as working memory, reasoning, and cognitive control. Studying these developmental reconfigurations in functional networks is vital for uncovering the neural mechanisms underlying cognitive maturation and for identifying potential deviations linked to neurodevelopmental disorders.

Resting-state functional magnetic resonance imaging (rs-fMRI) has emerged as a valuable tool for studying the neural basis of cognitive development (Cosío-Guirado et al., 2024). Graph theory analysis is widely used for analyzing rs-fMRI data (Rubinov & Sporns, 2010) by modeling the brain as a complex network in which nodes represent brain regions and edges represent their connectivity. Graph theory metrics quantify network properties based on these nodes and edges (e.g., degree measures number of connections and clustering coefficient measures local interconnectedness), offering insights into functional reconfiguration during development (Wang, 2010). Key topological properties of global functional brain architecture reach relative maturity by around age eight (Menon, 2013). However, existing literature suggests that global metrics (e.g., normalized mutual information, adjusted rand index) may lack the sensitivity to detect regional network alterations (Jalbrzikowski et al., 2019; Marek et al., 2015). Developmental reconfigurations nevertheless continue throughout late childhood and adolescence, with maturation occurring primarily at the regional level (Baker et al., 2015; López-Vicente et al., 2021). These findings indicate that region-level graph theory metrics (Brandes, 2001; Kleinberg, 1999; Latora & Marchiori, 2001) may be more sensitive for capturing these developmental processes.

In the last decade, increasing research has investigated the behavioral significance of developmental reconfiguration in functional networks. Specifically, increased functional connectivity (FC) between parietal-frontal brain regions is associated with higher nonverbal intelligence in children, with stronger associations observed in girls (Langeslag et al., 2013). Age-related cognitive development is associated with the segregation level of the sensorimotor and default mode network measured by between-network connectivity (Gu et al., 2015). Higher reading achievement is positively associated with FC between the dorsal striatum and the dorsal attention network (Jolles et al., 2020). Evidence has suggested that these brain-behavior relationships gradually change across development (Barber et al., 2013; Koyama et al., 2011; Wendelken et al., 2016), which underscores the importance of investigating developmental reconfiguration in relation to cognitive performance. Moreover, both FC between regions and cognitive processes have been found to differ by sex (Icer et al., 2020; Satterthwaite et al., 2015; Tomasi & Volkow, 2023). Although sex differences in the prevalence and presentation of psychiatric disorders have been well documented (Thibaut, 2016), the brain mechanisms driving the differences remain unclear. The majority of current studies examining developmental reconfiguration emphasized global FC or adult-derived functional networks. Investigating regional developmental trajectories and sex-specific patterns in typically developing children could clarify how network reorganization supports cognitive maturation. Given that cognitive impairments in various domains (e.g., sensory processing, attention, and facial recognition) are often considered as hallmark features of neurodevelopmental disorders, understanding normative brain network maturation provides the opportunity to identify deviations in maturation which may be markers of symptoms of impaired neural and cognitive development.

In the current study, we analyzed resting-state fMRI data from 528 adolescents and young adults aged 8–21 from the Human Connectome Project in Development (HCP-D) dataset. Our primary aim was to characterize how regional network properties evolve across adolescence and to examine whether these developmental trajectories differ between sexes.

To capture local aspects of functional reconfiguration, we focused on region-level graph theory metrics. Furthermore, we investigated how age-related changes in these regional metrics are associated with cognitive maturation.

## 2. Materials & Methods

### 2.1. Participants

We used the latest release of the HCP-D dataset (available at https://www.humanconnectome.org/), which extended the adult HCP study to examine brain development in healthy children acquired at four sites across the US. A detailed description of the dataset has been previously published (Somerville et al., 2018). In brief, the project recruited healthy individuals aged 5 to 21 years to capture a representative sample of children, adolescents, and young adults. Participants younger than 18 years were accompanied by a parent or legal guardian, who will provide written informed consent for the child’s participation. Participants were excluded if they (a) were unable to complete study procedures, (b) had medical conditions that could interfere with participation or compromise confidentiality, (c) had a history of neurodevelopmental disorders, or (d) presented contraindications for MRI. In the current release (version 2.0), baseline data were available for 625 participants. After excluding individuals with excessive head motion during scanning, as described in the following section, the final analytic sample included 528 participants.

### 2.2. Image acquisition and pre-processing

During HCP-D rs-fMRI acquisition, participants were instructed to stay still, stay awake, and blink normally while fixating on a central crosshair. For participants aged 8 years and older, 26 minutes of resting-state scanning were acquired across four runs over two consecutive days using the 3T Siemens Prisma platform (TR/TE= 800/37 ms, flip = 52, FOV= 208 × 180 mm, matrix = 104 × 90, slices = 72, voxel size = 2.0 × 2.0 × 2.0 mm). For younger participants (ages 5–7 years), each run lasted 3.5 minutes for a total of six runs (21 minutes of rs-fMRI data) (Harms et al., 2018).

Preprocessing followed the HCP minimal preprocessing pipeline (Glasser et al., 2013). Several covariates were regressed out, including linear and quadratic drift, mean cerebral-spinal fluid signal, mean white matter signal, and overall global signal. Images were then bandpass filtered between 0.008 Hz and 0.12 Hz. To mitigate the confounding effect of head motion, the mean framewise displacement (Power et al., 2012) was calculated for each run for each participant, and runs with a mean framewise displacement exceeding 0.3 mm were excluded. Additional preprocessing steps to minimize motion confounds were performed, including iterative smoothing, regression of 24 motion parameters (6 rigid-body parameters, 6 temporal derivatives of these parameters, and these 12 parameters squared), and frame censoring for volumes with displacement>0.3 mm. Participants whose rs-fMRI data contained more than 30% frames were excluded, resulting a final sample of 528 participants (age 67-263 months [5-21 years]; 290 females). We used the Glasser atlas (360 cortical regions) to parcellate the regions of interest (ROIs). For each participant, the mean time series of each ROI was extracted and ROI-to-ROI FC matrices were generated using Fisher□z-transformed Pearson correlations. Site effect were harmonized using NeuroCombat (Fortin et al., 2018).

### 2.3. Network construction and graph theory metrics

For each participant, the FC matrix was converted to absolute values and thresholded into a binary connectivity matrix (i.e., adjacency matrix) which only retains the strongest 1% of connections. Specifically, the threshold value was computed as the 99th percentile of edge weights, excluding the diagonal (i.e., 1% threshold). To assess robustness, PARAFAC analyses were repeated at 5% and 10% density thresholds. The entry of the adjacency matrix was set to 0 if the edge weight between a pair of regions was below the threshold and 1 if otherwise. Each adjacency matrix was visualized as a network graph constituted by a number of nodes (i.e., regions) and a number of edges. For each participant, three region graph theory metrics (betweenness centrality, hub score, and local efficiency) were computed for each brain region to characterize functional integration and segregation during development (Fair et al., 2008). **Fig. 1** illustrates the schematic representation of the analysis process.

**Fig. 1.**
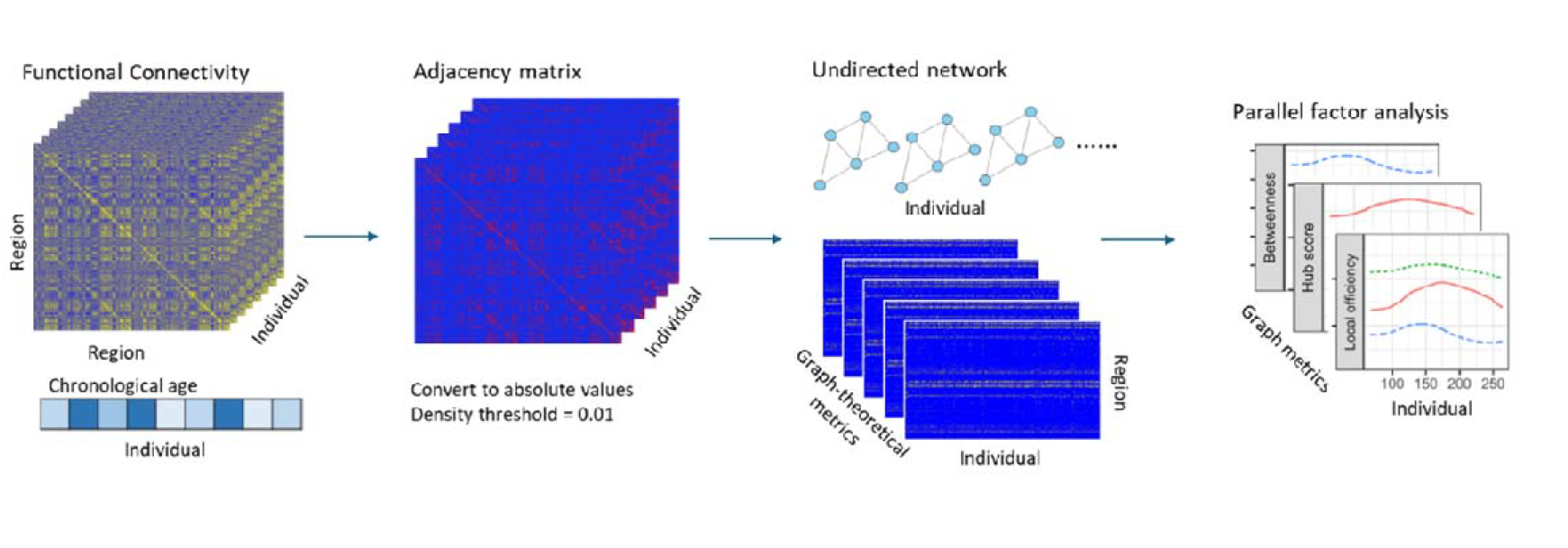
Schematic representation of the analysis process

#### 2.3.1. Functional integration measures

Functional integration refers to the brain’s capacity to efficiently coordinate and combine information across distributed specialized regions (Rubinov & Sporns, 2010).

*Betweenness centrality:* quantifies network integration by estimating the long-range centrality of each node based on the fraction of shortest paths between all other nodes that pass through it (Brandes, 2001). A higher betweenness centrality value of a given node (in this case, brain region) signifies a more critical role for that node in facilitating the communication between networks.

*Hub score*: quantifies a region’s local centrality by considering its direct connections, reflecting both the number and significance of these links (Kleinberg, 1999). Hubs refer to important brain regions that interact with many other regions to facilitate communications within and between networks. An higher hub score indicates that a region occupies a more central or influential position within the network.

#### 2.3.2. Functional segregation measure

Functional segregation in the brain is the ability to support specialized processing to occur within densely interconnected groups of brain regions.

*Local efficiency:* measures the average inverse path length between a region’s neighbors, reflecting the integration specifically within a node’s local environment (Latora & Marchiori, 2001). For example, decreased local efficiency in a brain region may indicate impaired specialized processing within that functional network.

### 2.4. Parallel factor analysis

In the present study, we applied parallel factor analysis (PARAFAC) to identify latent patterns in age-related changes of graph theory metrics. Specifically, we first constructed a three-way array with dimensions 528 age points × 360 regions × 3 metrics, denoting by *X*= {*x*_*ijk*_}_*I*×*J*×*K*_. Each entry *x*_*ijk*_ represents the i-th age point (from the *i*-th individual) of *j*-th region for the *k*-th graph theory metric. Age was measured in months. To avoid ties in age (i.e., participants with the same age value), random number generation from [0,1) was added to age, thus each participant represents one point on the continuous chronological age scale. We then decomposed the array using PARAFAC into sets of scores and loadings to describe the data in a more condensed and interpretable form. As a generalization of bilinear principal component analysis for multi-way data, PARAFAC simultaneously fits multiple ‘slices’ of a multi-way array in terms of a common set of factors with differing relative weights in each array (Harshman & Lundy, 1994). Each graph theory metric was scaled by dividing its standard deviation to preserve the non-negativity during model fitting process, enabling meaningful interpretation at factor level. Compared to other multi-way methods like three-mode principal component analysis Tucker3, PARAFAC is simpler and more restricted but generates a robust and interpretable model (Bro, 1997).

Following previous studies, we estimated PARAFAC weights using an alternating least squares algorithm that minimizes the sum of squared errors, with a non-negative spline constraint for the age dimension to encourage a smooth trajectory and a non-negative orthogonal constraint for the region dimension to ensure interpretability (Harshman & Lundy, 1994; Helwig, 2017). For the non-negative spline constraint, we determined the number of knots (i.e., points that join polynomial segments) for the spline using ordinary cross-validation and the same number of knots was set for all three metrics. The number of factors (i.e., principal components characterizing the three-way arrays) was determined using the Consistency Diagnostic metric (CORCO) (Bro & Kiers, 2003). We repeated the analytical pipeline separately for males (n=238) and females (n=290) to better understand sex-specific functional reconfiguration across development. Post hoc analyses were conducted to quantify curve similarity between sex-specific patterns and pattern of entire samples using the L2 distance.

All regions were modeled independently in the PARAFAC analysis, without being constrained by their assignment to adult-derived functional networks (e.g., default mode network, visual network, somatomotor network) defined in standard brain-parcellation schemes. Then, we assigned each region to functional networks according to Cole-Anticevic cortical Network Partition (version 1.1), which includes 12 predefined networks derived from adult samples, as presented in Supplementary Fig. S1 (Ji et al., 2019). This allowed us to assess whether developmental patterns were consistent with or distinct from those of other regions within the same network. By examining the factor weights along the age dimension for each metric, we identified characteristic developmental trajectories of local network properties within each sample.

### 2.5. Cognitive performance in age-specific developmental subgroups

We examined the associations between factor weights at the metric dimension for each participant (i.e., averaged metric values across all regions for each participant) and cognitive measures to elucidate how local network reconfigurations correspond to cognitive development during this period. Cognitive performance were measured by NIH Toolbox Cognition Battery, which is a battery of seven cognitive tests included two measures of crystallized abilities (Picture Vocabulary Test and the Oral Reading Recognition Test) and six measures of fluid abilities (the Toolbox Dimensional Change Card Sort (DCCS) Test, the Toolbox Flanker Inhibitory Control and Attention Test, the Toolbox Picture Sequence Memory Test, the Toolbox List Sorting Working Memory Test, and the Toolbox Pattern Comparison Processing Speed Test) (Akshoomoff et al., 2013; Gershon et al., 2010; Weintraub et al., 2013). Crystallized cognition composite, fluid cognition composite, and cognitive function composite scores were calculated by averaging the scaled scores from these tests. Additionally, an early childhood composite score is derived from four cognitive tests of Early Childhood Battery, including Picture Vocabulary, Flanker, DCCS, and Picture Sequence Memory (Weintraub et al., 2013). Partial correlation controlled for age was calculated between the four cognition composite scores and the average factor weights of each metric in three developmental groups: 1) age below 156 months (<13 years), 2) age between 156 and 216 months (13-18 years), and 3) age above 216 months (>18 years). The same analyses were repeated in female and male samples.

## 3. Results

### 3.1. Parallel factor analysis

Sample characteristics of the study cohort are presented in **Table 1**. The PARAFAC analysis identified three factors. **Fig. 2** illustrates the weights of each brain region on each factor in full, female, and male samples. Overall, the factor weight distribution remained consistent between the sexes, though minor differences appeared in the CAB-NP functional networks involved in each factor. Factor 1 showed strong weights predominantly from regions of visual networks. Factor 2 demonstrated broader network engagement, with regions of multiple networks showing moderate loadings distributed across diverse cortical areas. Factor 3 revealed increased contributions from regions of higher-order networks, notably the frontoparietal, dorsal-attention, and cingulo-opercular networks. Accordingly, we refer to factor 1 as the *visual factor*, factor 2 as the *multimodal integration factor*, and factor 3 as the *higher-order factor*, reflecting network affiliations of their predominant region loadings. The corresponding factor weights of regions from specific networks are present in Fig. S2.

**Table 1.** Sample characteristics.

| Characteristic | All<br>N = 528 | Female<br>N = 290 | Male<br>N = 238 |
| --- | --- | --- | --- |
| Age in months, Mean (SD) | 179 (47.5) | 177 (48.6) | 181 (46.2) |
| Race, n (%)<sup>1</sup> |  |  |  |
| White | 330 (62.5%) | 183 (63.1%) | 147 (61.8%) |
| Non-white | 185 (35.0%) | 98 (33.8%) | 87 (36.6%) |
| Ethnic group, n (%)<sup>2</sup> |  |  |  |
| Hispanic or Latino | 79 (15.0%) | 45 (15.5%) | 34 (14.3%) |
| Not Hispanic or Latino | 440 (83.3%) | 238 (82.1%) | 202 (84.9%) |
| Cognitive performance<sup>3</sup> | 104.7 (13.0) | 105.1 (12.5) | 104.3 (13.7) |

<sup>1</sup> There are 13 with unknown race, including 9 females and 4 males
<sup>2</sup> There are 9 with unknown ethnic groups, including 7 females and 2 males
<sup>3</sup> NIH toolbox total cognition composite score missing for 154, including 88 females and 66 males

**Fig. 2.**
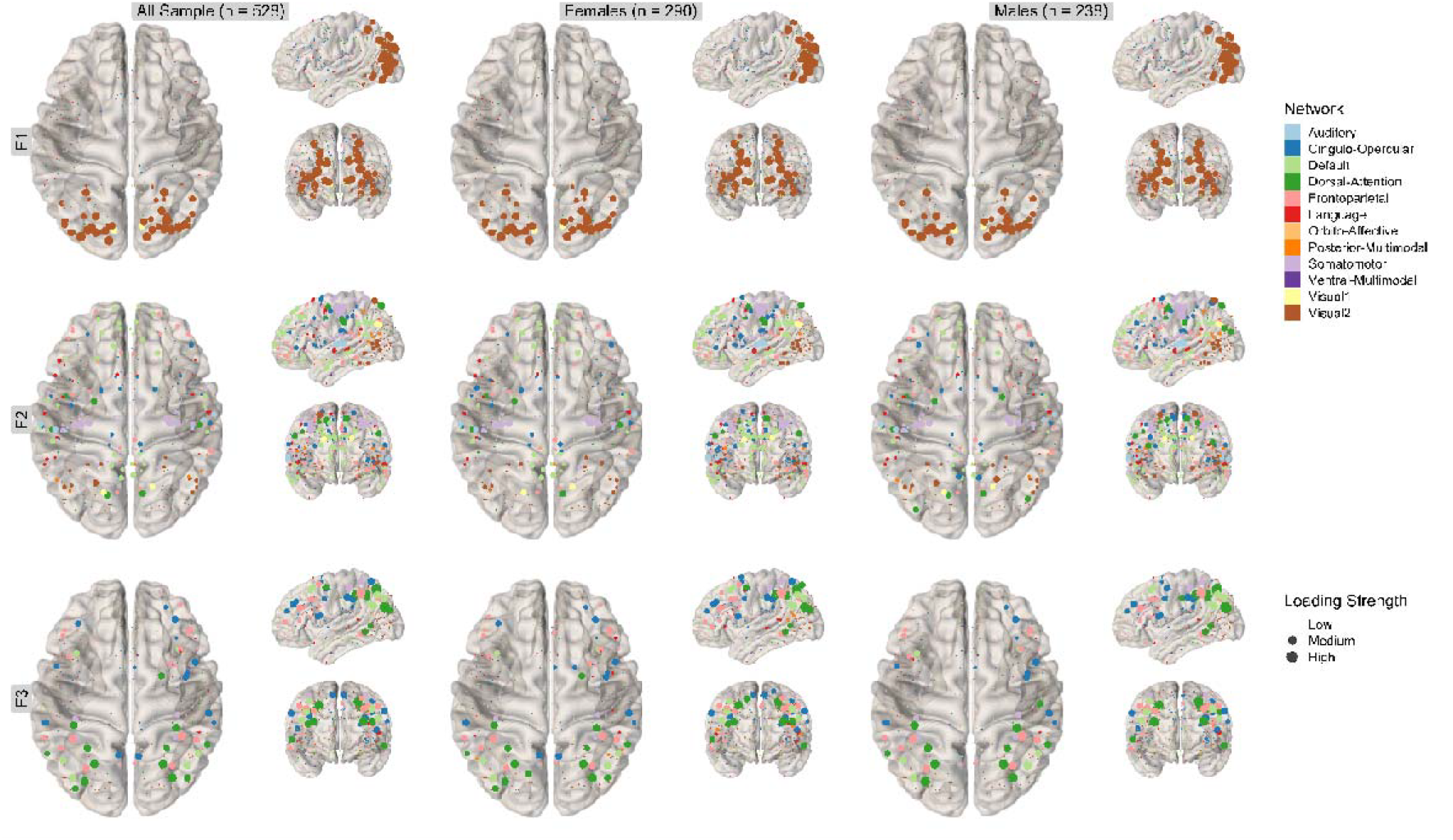
Factor weights of each brain region on each factor in entire sample, females, and males. F1: visual factor; F2: multimodal integration factor; F3: higher-order factor

**Fig. 3.**
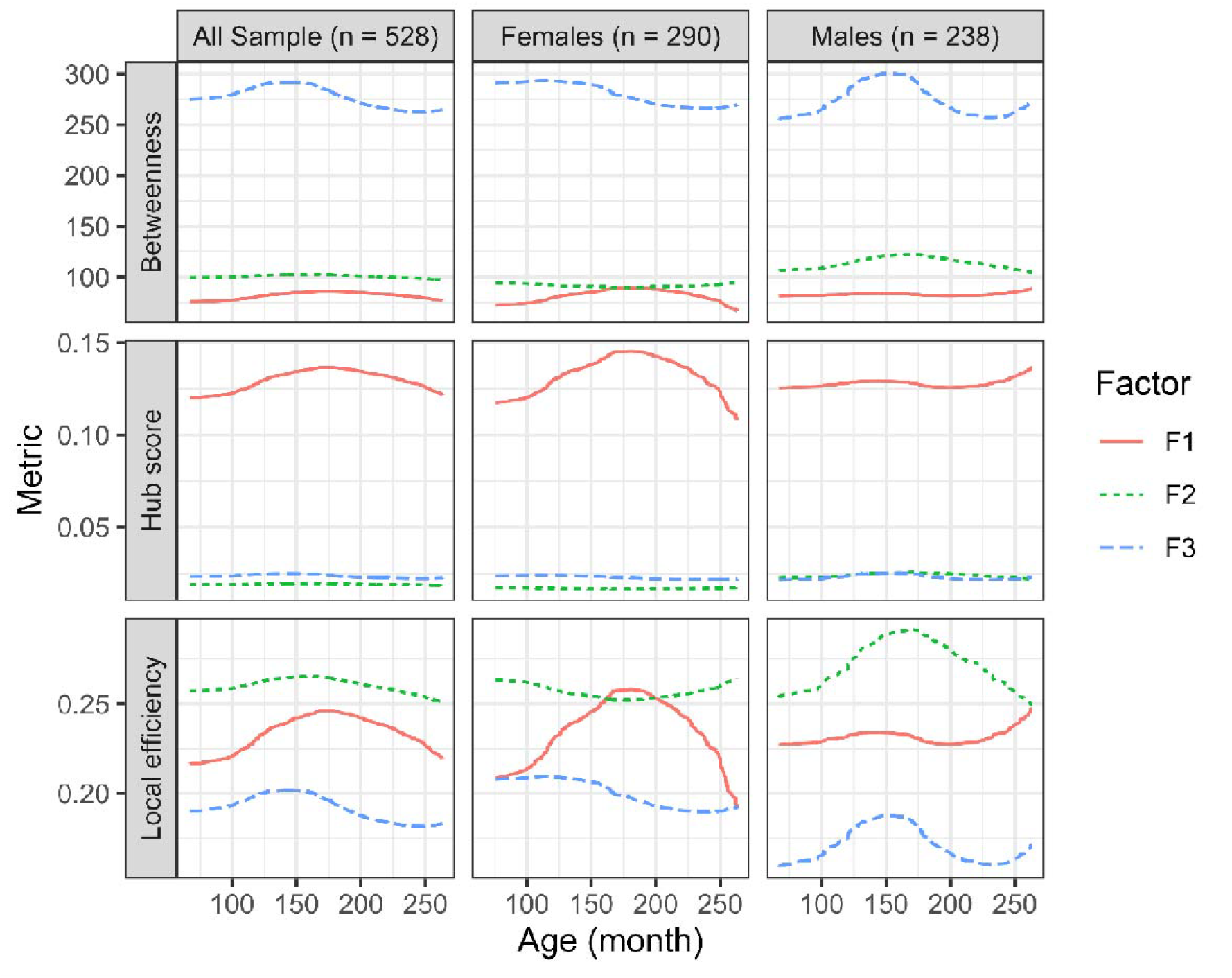
Average factor weights of all regions along the age dimension in entire sample, females, and males. F1: visual factor; F2: multimodal integration factor; F3: higher-order factor

In **Fig. 3**, we plot average factor weights of all regions along the age dimension to reveal the variation pattern of the three factors in each metric. Across the entire sample, betweenness centrality and hub score remained relatively stable over development, with direction changes observed for betweenness centrality at month 139 (11.5 years) for the higher-order factor, and hub score at month 175 (14.6 years) for the visual factor. In contrast, the local efficiency of all factors exhibited direction changes at or after 139 months (11.5 years).

**Fig. 3.**
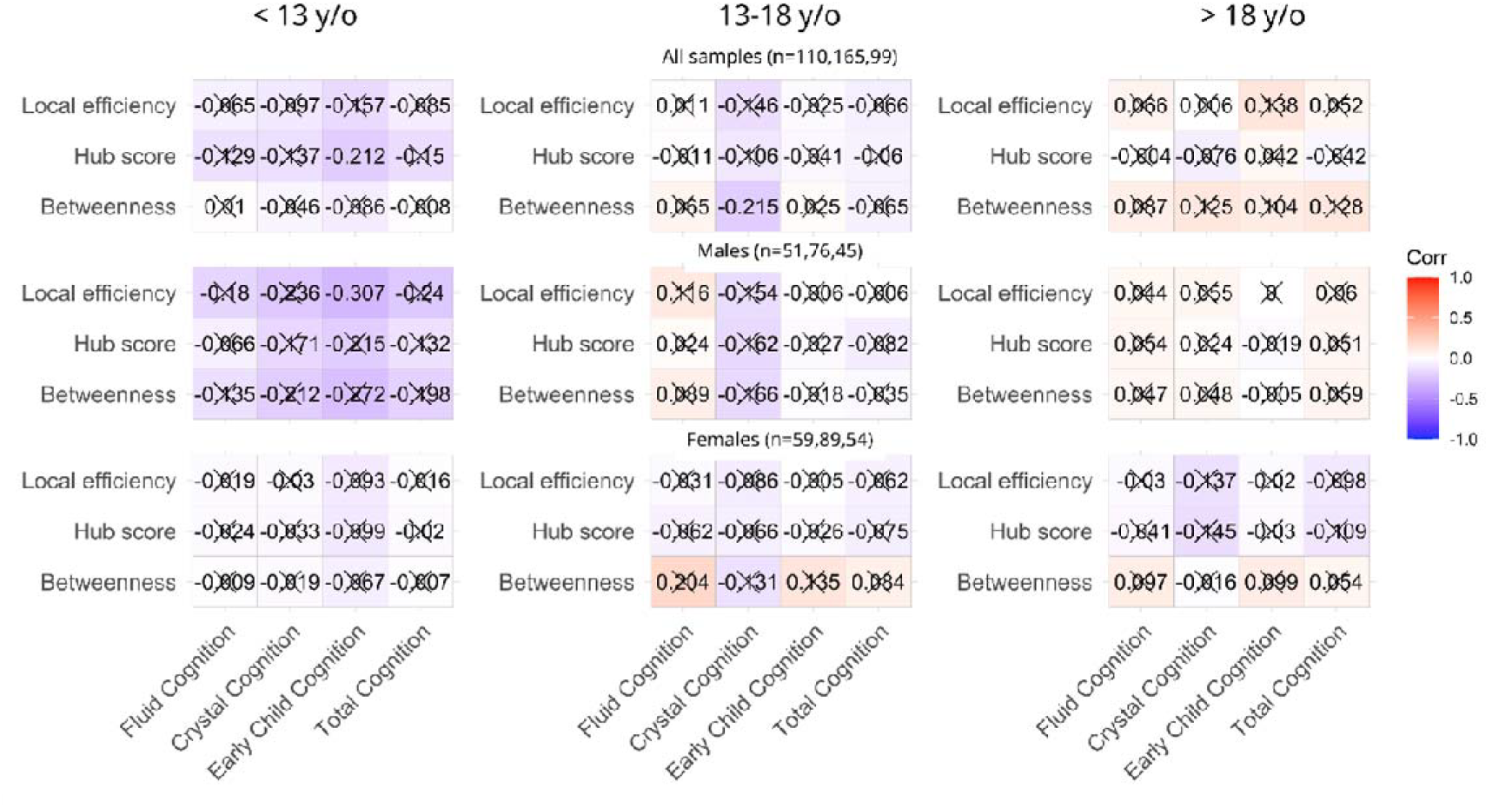
Partial correlations between cognitive performance scores and average weights of graph theory metrics in age subgroups

### 3.2. Sex-specific functional reconfiguration over development

We found notable sex-specific variation in functional reconfiguration patterns across the various age subgroups. In females, hub score and local efficiency peaked around month 175 in the visual factor, whereas betweenness centrality and local efficiency decreased with age in higher-order factor. Conversely, in males, all three metrics showed an upward trend in visual factor, whereas betweenness centrality and local efficiency changed direction at month 150 in higher-order factor.

Overall, factor patterns differed by sex across the three metrics. The multimodal integration factor and higher-order factor were generally stable with no directional changes in females across all regional graph theory metrics, while peaks were observed for males in betweenness and local efficiency of these factors. The visual factor showed the opposite for males and females. For the multimodal integration factor, the entire-sample pattern more closely resembled that of males (L2 distance: d=8.40) than females (d=15.23). The opposite pattern was observed for the visual factor (males: d=3.80; females: d=3.02) and higher-order factor (males: d=8.98; females: d=7.86), where the entire-sample pattern showed greater similarity to the female pattern.

### 3.3. Age-specific association between metrics and cognitive performance

As shown in **Fig. 4**, the early childhood cognitive composite score was negatively correlated with hub score in younger children (full sample, <13 years old; r = -0.212, p = 0.027) and with local efficiency in younger males (r = -0.215, p = 0.006). Crystal cognition composite score was negatively correlated with betweenness centrality in older children (13-18 years old, r = -0.307, p = 0.030). There were no other significant correlations between cognitive performance scores and average weights of graph theory metrics. The residual-by-residual plots corresponding to the partial correlations between metrics and cognitive scores were included in the supplementary materials (Fig. S3-S5).

## 4. Discussion

This study utilized graph theory metrics to characterize regional functional reconfiguration in typically developing children and youth. Our findings demonstrate the nonlinear nature of functional network reconfiguration across development, illustrated by the age-related patterns of the multimodal integration and higher-order factors in males and the visual factor in females, which reflect dynamic brain maturation through concurrent pruning and rewiring processes rather than linear or unidirectional trajectories (Menon, 2013). We found that the spatial distribution of regions’ factor weights was generally consistent across sexes, aligning with previous work suggesting key topological features of global functional brain architecture are established at an early age (Gao et al., 2015). However, sex-specific differences were observed in the timing and nonlinearity of network reconfiguration related to the formulation of higher-order cognition. Specifically, compared to males, females exhibited earlier peaks and an earlier decline in higher-order networks metrics (betweenness centrality and local efficiency). This pattern suggests an earlier functional maturation in females, reflected in the dynamic integration and segregation of networks supporting higher-order cognitive functions. Stronger inverted U-shaped trends of integration and segregation were found for the visual factor in females, whereas this trend was observed in higher-order factor and multimodal integration factor in males. These findings suggest sex-specific differences presented in the reconfiguration of regions that support higher-order cognitive function and the non-linear developmental trends exhibited in different regions across sex. Finally, age- and sex-specific effects were found between graph theory metrics and cognitive performance scores.

### 4.1. Sex-specific difference in developmental reconfiguration

Males showed greater variability than females in the higher-order factor and multimodal integration factor but not visual factor, which is consistent with prior findings of selective greater male-than-female variability in specific FC measures, including heteroscedastic changes across cortical and subcortical networks (Bottenhorn et al., 2023). This pattern also aligns with an Italian population-based study of over 26 million children showing greater male variability in mathematics but not reading performance (Esposito et al., 2025). Together with exisitng evidence, our findings provides evidence to support that the greater male variability hypothesis (Ellis, 1911; Forde et al., 2020) is selective, applying primarily to specific features rather than ubiquitously across all human characteristics. In contrast, the higher local efficiency (i.e., more segregated networks) in females within the higher-order factor suggests that higher-order brain networks mature earlier and are more stable throughout development in females. These findings are consistent with previous studies suggesting that females experience earlier maturation of functional networks than males, as reflected by greater segregation and specialization of higher-order systems (Grayson & Fair, 2017; Menon, 2013).

Extensive evidence supports that primary sensory networks mature by approximately age two and show minimal developmental change thereafter (Gao et al., 2015; Gu et al., 2015; Homae et al., 2010). However, our results revealed apparent age-related changes in the development of visual regions in females (invert U-shape across all metrics). Our findings may suggest that specific visual regions which support complex cognitive skills, such as reading, continue to undergo subtle reconfiguration to advance cognitive processing (Grayson & Fair, 2017). Consistent with our findings, prior work has shown that girls develop stronger interhemispheric connections between visual orthographic regions and other reading-related areas with age (Liang et al., 2022).

### 4.2. Age-specific relationship between metrics and cognition

We did not observe significant sex or age effects on the partial correlations between graph-theory metrics and cognitive function across the entire sample. Similar null findings have been reported by previous studies investigating intelligence-related differences in summary global parameters after controlling for sex (Wu et al., 2013). However, within the subgroup of males younger than 13 years, local efficiency was negatively associated with early childhood cognitive ability, an effect not observed among females of the same age range. Local efficiency, reflecting the capacity for functional segregation that supports specialized processing to facilitate cognitive performance, is particularly relevant for early childhood cognition. This sex-specific pattern may reflect males’ relatively protracted functional maturationmay reflect the relatively protracted functional maturation characteristic of males. During early adolescence, this maturational lag may transiently stronger brain–behavior couplings prior to the stabilization of large-scale network reconfiguration. Overall, our results reveal the role of functional network topology in supporting cognition in typically developing individuals, highlighting subtle, age- and sex-specific variations that warrant further longitudinal investigation.

### 4.3. Strengths and limitations

A key strength of this study lies in its exclusive focus on typically developing youth, enabling a detailed characterization of normative brain network maturation. By characterizing normative patterns of functional network reconfiguration, these findings lay the groundwork for evaluating how deviations of typical functional development may underlie psychiatric symptoms. Despite providing valuable insights into functional network reconfiguration, this study has several important limitations. First, the use of cross-sectional rather than longitudinal data limits the ability to examine individual-specific developmental trajectories and causal inferences regarding network reconfiguration over time. Second, the use of a particular threshold in network construction would largely affect the density of a network. To address potential bias from network thresholding, we repeated the analysis, retaining only the strongest 5% or 10% of connections (Supplementary Figs. S6–S7). Both thresholds yielded a two-factor solution, in which regions from the visual and higher-order factors identified in the primary analysis merged into a single factor, likely reflecting increased factor saturation at higher network densities, where weaker connections obscure factor boundaries. When a three-factor solution was forced, however, factor loadings remained generally consistent with those from the primary analysis (Supplementary Figs. S8–S9). These findings suggest that denser networks capture more distributed connectivity patterns, whereas sparser networks emphasize core connectivity architecture. Our primary analysis used a 1% threshold to focus on the strongest functional connections, thereby maximizing signal-to-noise ratio and minimizing spurious correlations from weak or noisy edges. Future studies should systematically evaluate a range of thresholds to identify the optimal balance between network density and interpretability. Third, the lack of association between graph theory metrics and cognitive performance is likely due to the generally healthy individuals included in this cohort; thus, it cannot be generalized to other populations. These limitations highlight the need for future longitudinal, multi-cohort studies to deepen understanding of functional network development.

## 5. Conclusion

In conclusion, this study demonstrates that normative neurodevelopment is driven by regionally specific reconfigurations of functional brain networks that global metrics alone are insufficient to detect, and that these subtle regional shifts may account for meaningful variance in cognitive performance. Critically, these trajectories diverge by sex, highlighting the necessity of sex-stratified analyses for capturing the true complexity of functional maturation. Together, these findings provide empirical support for theoretical accounts positing that cognitive maturation emerges from regional network reconfiguration. Future clinical investigations may build on this foundation by examining how deviations from these normative trajectories contribute to vulnerability for neurodevelopmental and psychiatric conditions.

## Acknowledgement

The current study is partially supported by R01MH124106. Data were obtained from the NIMH Data Archive (NDA Study 3221, doi:10.15154/bxfa-z706).

## Declaration of generative AI use

During the preparation of this work, the author(s) used Perplexity (Perplexity AI, San Francisco, CA) in order to check and improve English grammar. After using this tool, the author(s) reviewed and edited the content as needed and take(s) full responsibility for the content of the published article.

## Author Contributions

S.L. and C.Z.F. designed and conceptualized the study. C.Z.F. conducted statistical analyses. C.Z.F., H.N., and S.L. interpreted the study data. C.Z.F. wrote the first draft of the manuscript. C.Z.F. and H.N. made revisions with critical input from S.L., A.Z., and X.M. C.Z.F. and H.N. finalized the manuscript. All authors provided critical feedback to the manuscript and have approved the final manuscript.

