## Supplementary materials for "Regional reconfiguration of functional brain networks during childhood and adolescence: evaluating age and sex effect"

**Supplementary Fig. S1.** Cole-Anticevic cortical Network Partition of adult-derived functional network **Supplementary Fig. S2**. Regional factor weights with corresponding adult-derived functional network affiliations

**Supplementary Fig. S3**. Residual-by-residual plots corresponding to the partial correlations between metrics and cognitive scores in the entire sample

**Supplementary Fig. S4**. Residual-by-residual plots corresponding to the partial correlations between metrics and cognitive scores in males

**Supplementary Fig. S5**. Residual-by-residual plots corresponding to the partial correlations between metrics and cognitive scores in females

**Supplementary Fig. S6**. Factor loadings from parallel factor analysis of adjacency matrix retaining only the strongest 5% of connections

**Supplementary Fig. S7.** Factor loadings from parallel factor analysis of adjacency matrix retaining only the strongest 10% of connections

**Supplementary Fig. S8**. Factor loadings from parallel factor analysis of adjacency matrix retaining only the strongest 5% of connections (three-factor solution)

**Supplementary Fig. S9.** Factor loadings from parallel factor analysis of adjacency matrix retaining only the strongest 10% of connections (three-factor solution)

Fig. S1. Cole-Anticevic cortical Network Partition of adult-derived functional network

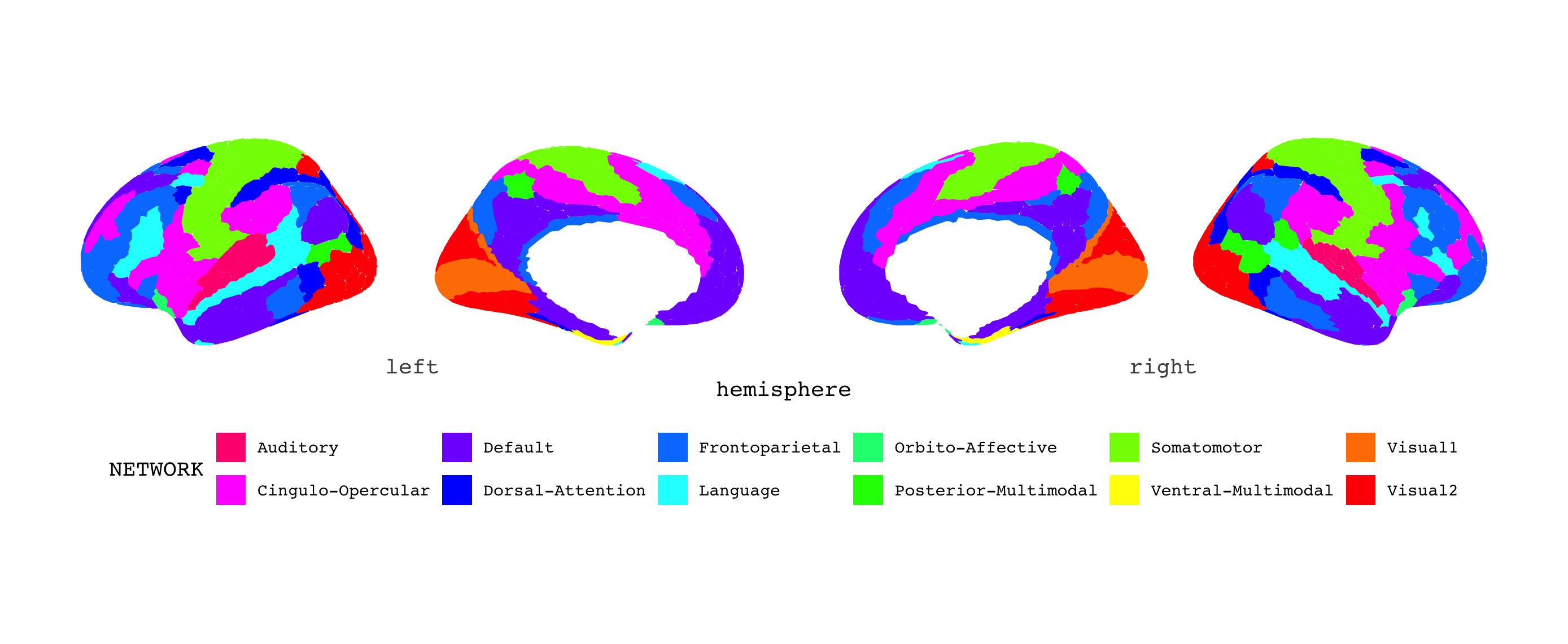

Fig. S2. Regional factor weights with corresponding adult-derived functional network affiliations

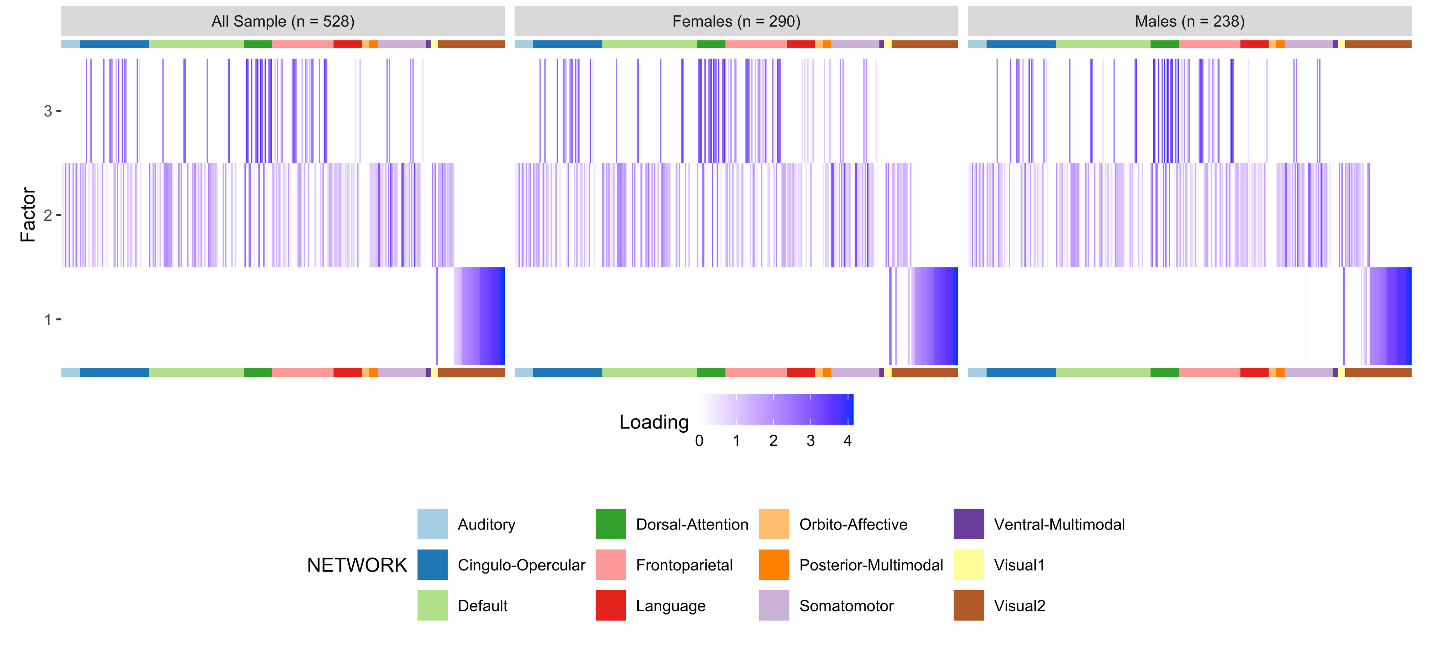

Fig. S3. Residual-by-residual plots corresponding to the partial correlations between metrics and cognitive scores in the entire sample

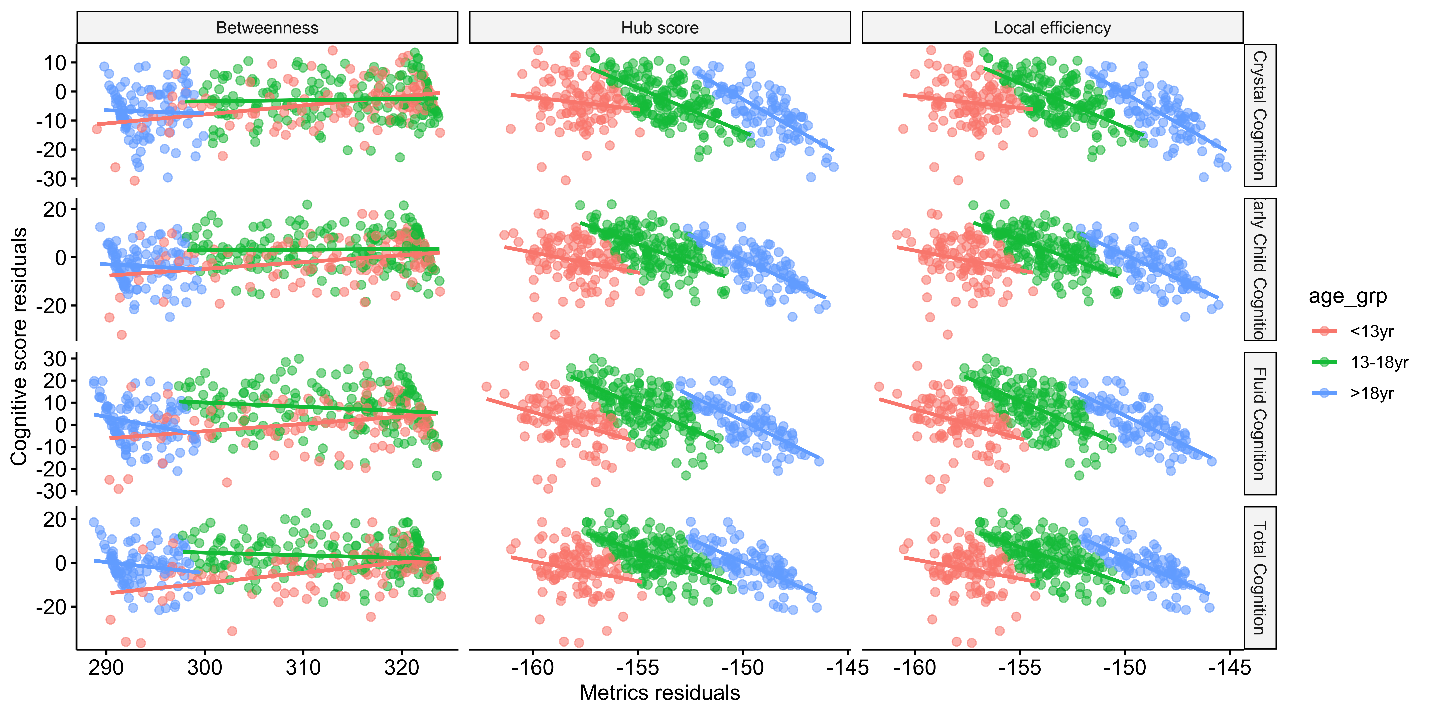

Fig. S4. Residual-by-residual plots corresponding to the partial correlations between metrics and cognitive scores in males

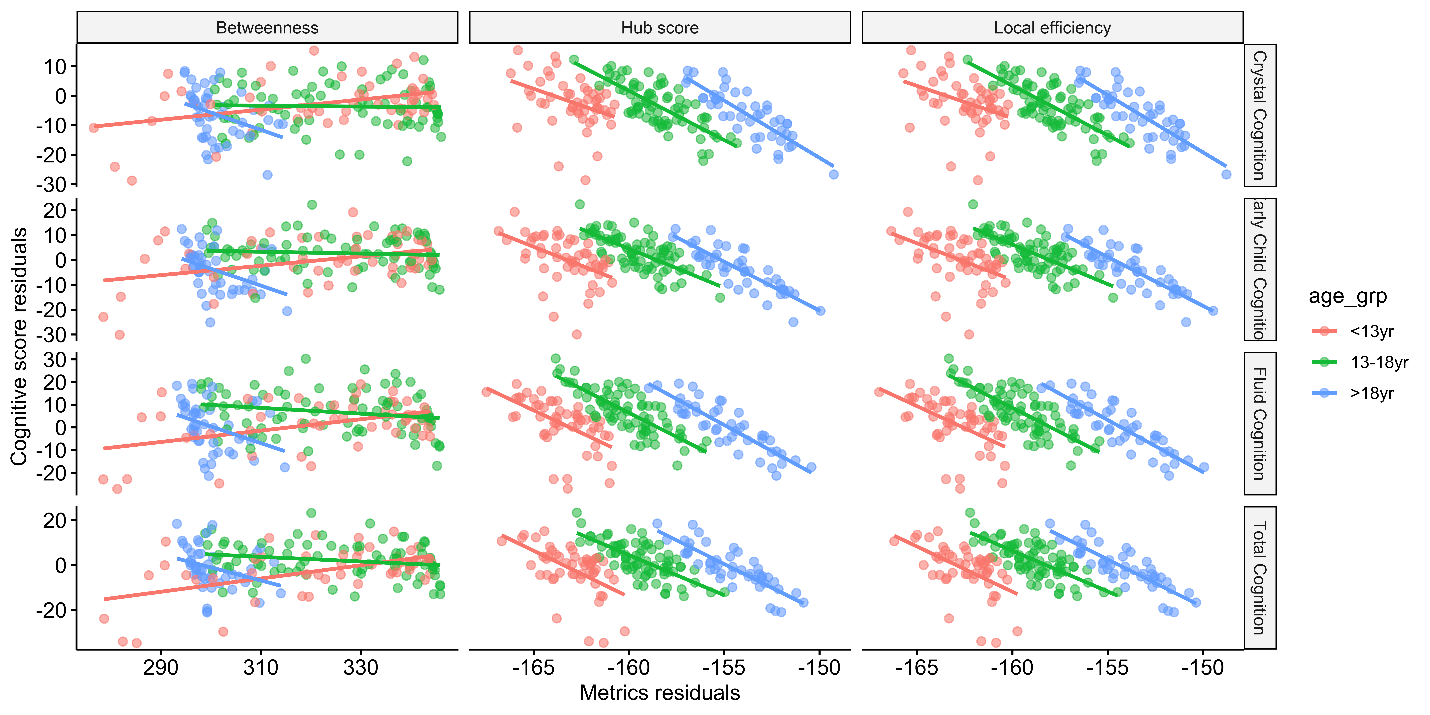

Fig. S5. Residual-by-residual plots corresponding to the partial correlations between metrics and cognitive scores in females

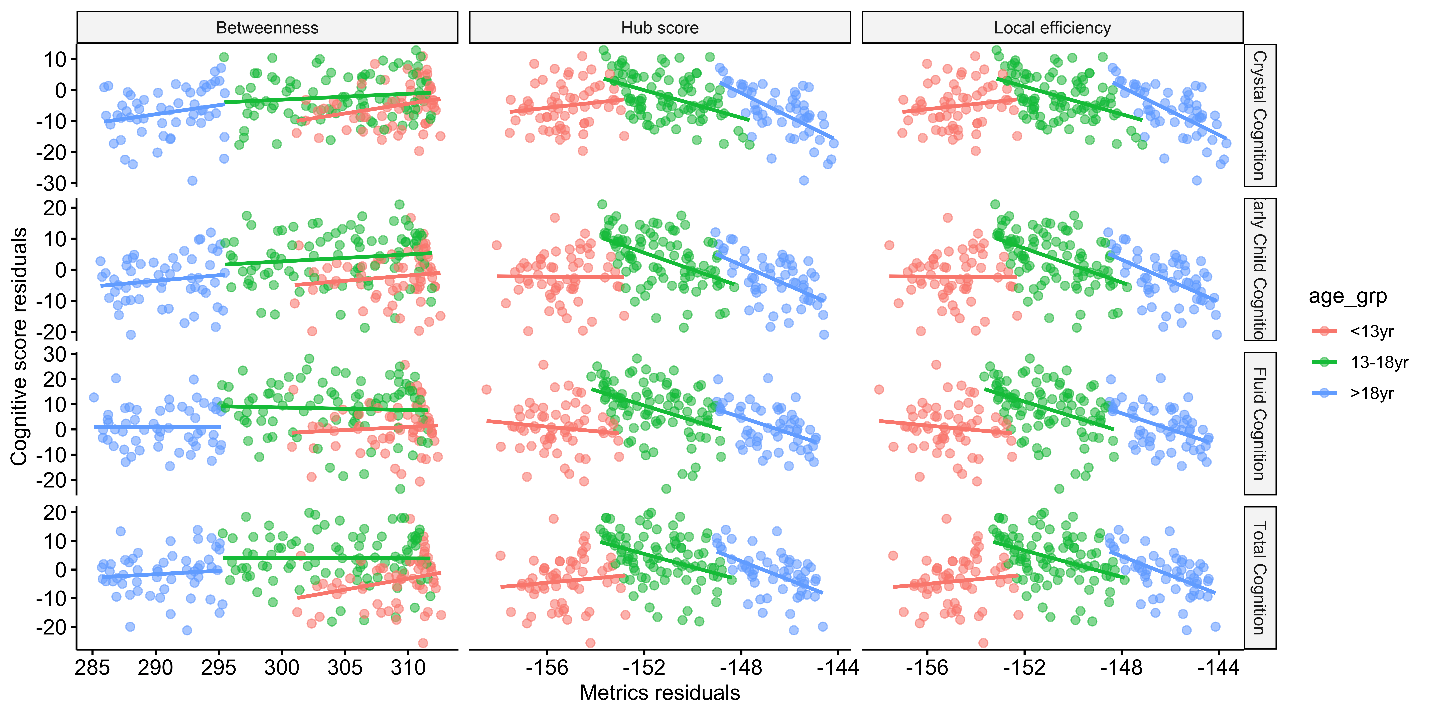

Fig. S6. Factor loadings from parallel factor analysis of adjacency matrix retaining only the strongest 5% of connections

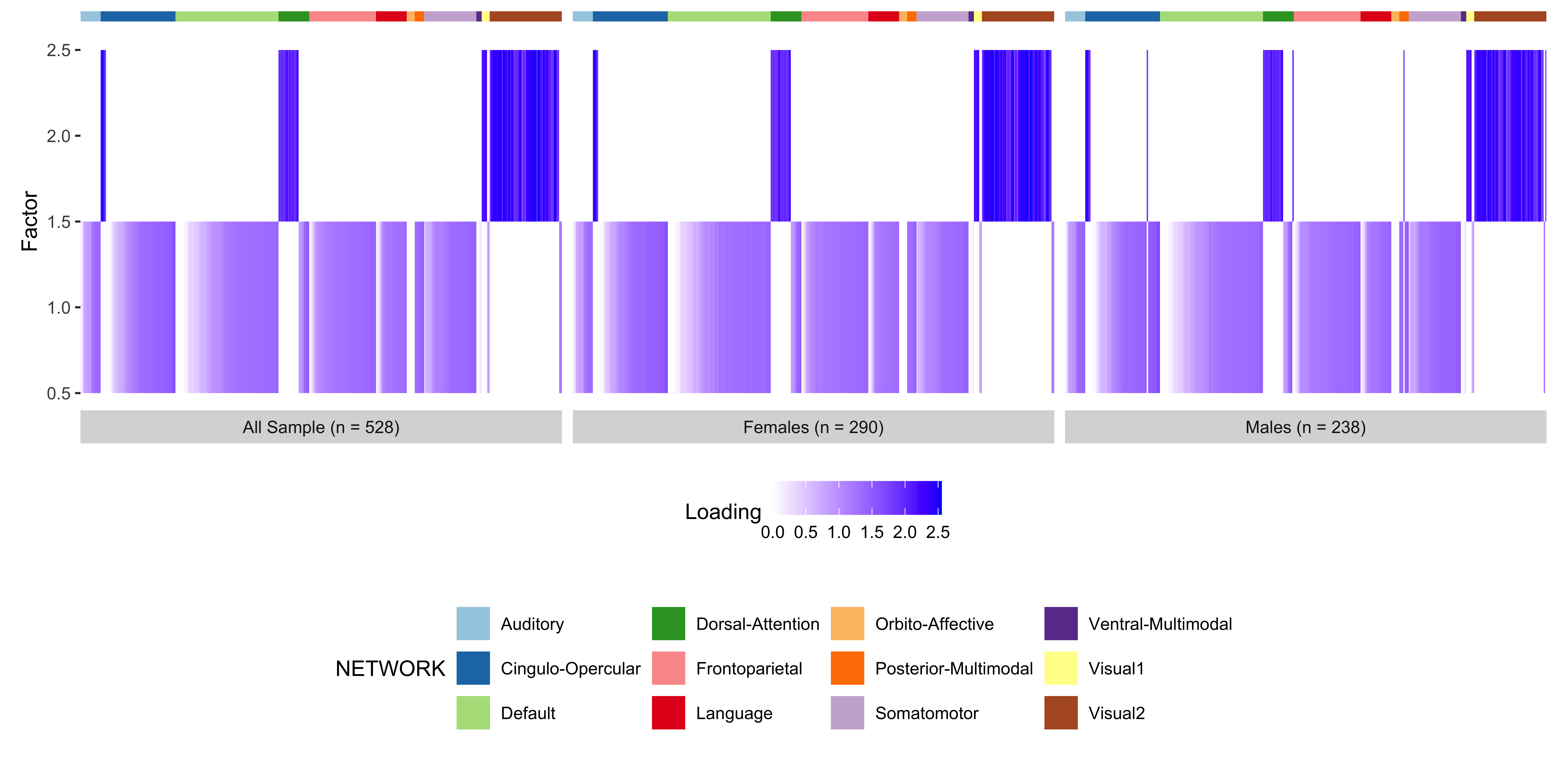

Fig. S7. Factor loadings from parallel factor analysis of adjacency matrix retaining only the strongest 10% of connections

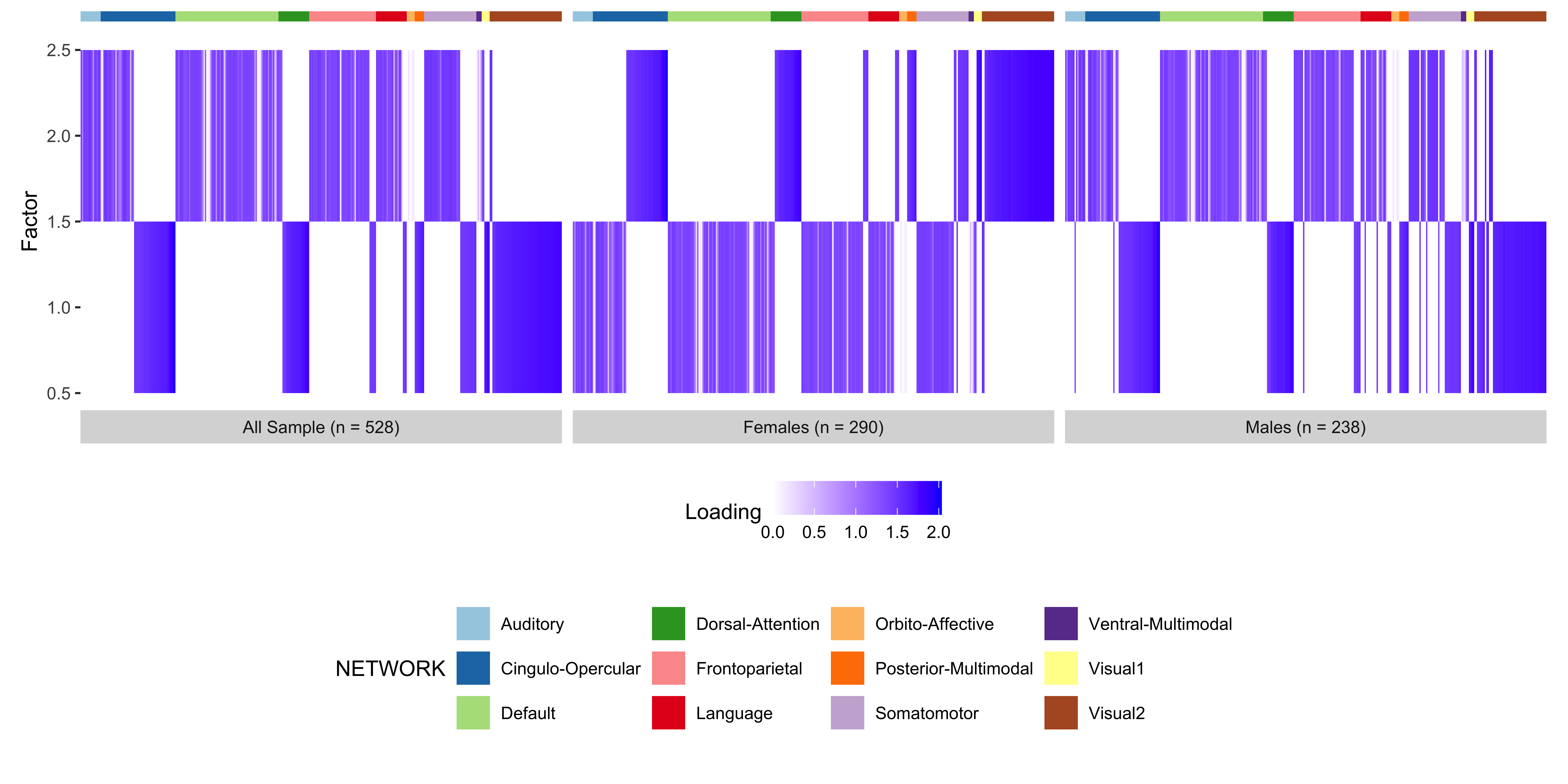

Fig. S8. Factor loadings from parallel factor analysis of adjacency matrix retaining only the strongest 5% of connections (three-factor solution)

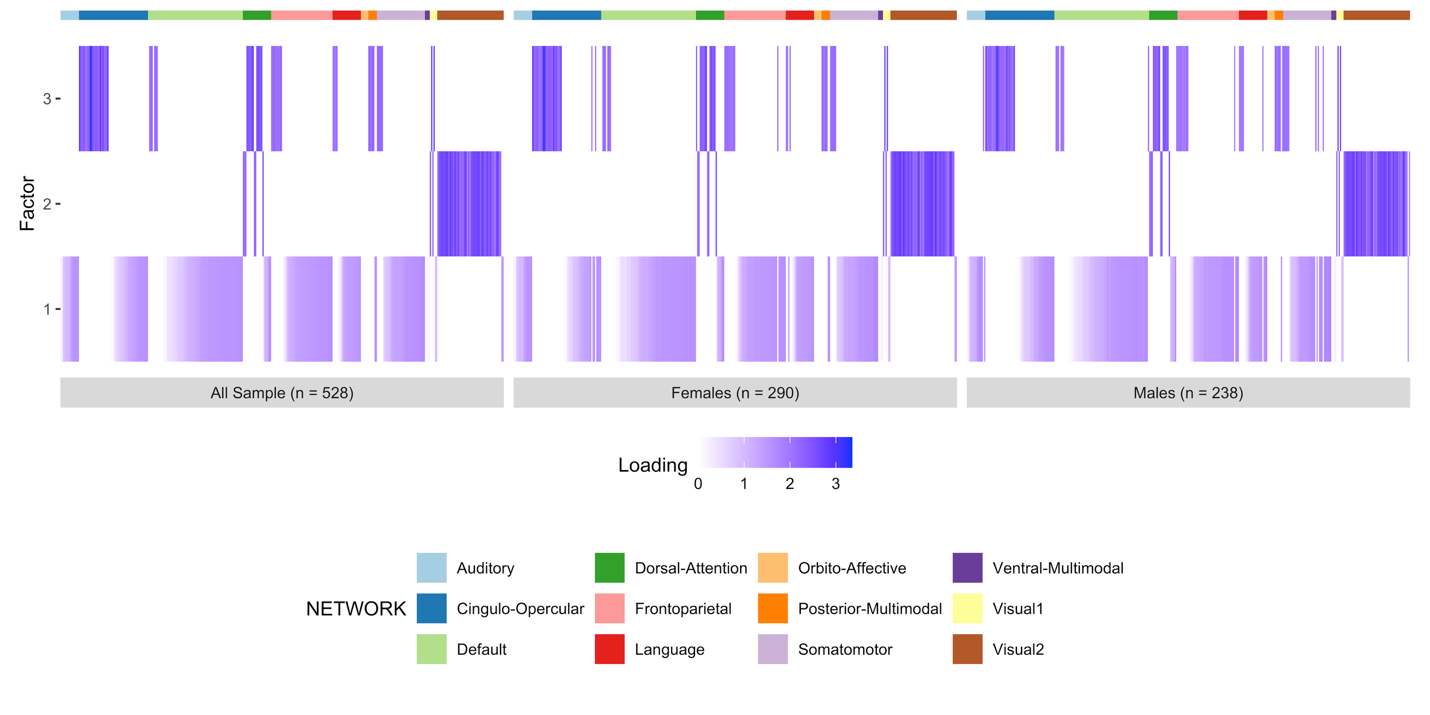

Fig. S9. Factor loadings from parallel factor analysis of adjacency matrix retaining only the strongest 10% of connections (three-factor solution)

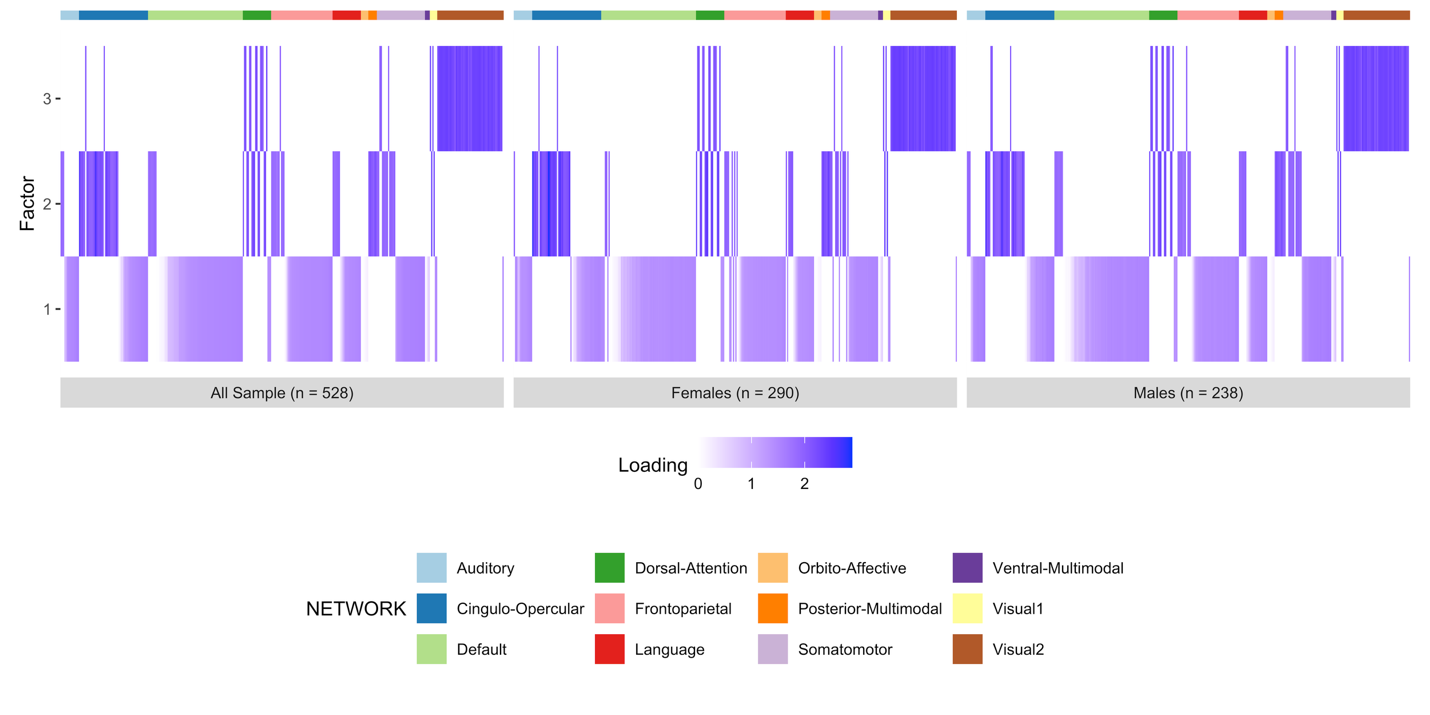
